# NON-TOXIC ACID-FREE GLYOXAL FIXATIVE FOR VETERINARY HISTOPATHOLOGY, IMMUNOHISTOCHEMISTRY AND MOLECULAR ANALYSIS

**DOI:** 10.1101/2023.05.05.539541

**Authors:** Valentina Zappulli, Filippo Torrigiani, Valentina Moccia, Paolo Detillo, Cecilia Gola, Lucia Minoli, Emanuela M. Morello, Erica I. Ferraris, Antonella Rigillo, Federico Caicci, Giulia Dalla Rovere, Davide De Biase, Lorenzo Riccio, Marco Rondena, Selina Iussich, Benedetta Bussolati

## Abstract

Formaldehyde fixation is worldwide the most used system for histopathological examination. However, its toxicity is well known, and preservation of proteins and nucleic acids is not optimal. Alternative fixatives warranting similar morphological quality of tissues and costs, but lacking toxicity and allowing better preservation of proteins and nucleic acids would therefore increase both safety of operators and quality of molecular analysis in pathology.

This multi-institutional study aimed to compare the morphological, histochemical, immunohistochemical (IHC), and molecular analyses outcomes of a newly patented, non-toxic, acid-free Glyoxal (GAF) fixative with neutral buffered formaldehyde (NBF). Tissues from a total of 73 subjects were analyzed, including 13 necropsies.

Gross features were preserved after GAF fixation, with no tissue hardening or discoloration. Cellular ultrastructure was also better preserved with GAF and histology and histochemistry on GAF-fixed samples showed good results when compared to NBF-fixed samples, with the exception of loss of tinctorial affinity of erythrocytes and mast cell granules. IHC analyses also showed comparable results with only slight and rare protocol adjustment. DNA and RNA yields were higher from GAF-fixed samples (P<0.05) and the tested genes (*p53* and *COX1*) were better amplified. RNA scope showed positive results for *c-KIT* expression in GAF-fixed mast cell tumors.

Based on these data, the non-toxic GAF fixative allows good macroscopical, histological and immunohistochemical analyses of tissue samples, including on-field application, and better molecular analyses when compared to NBF. This represents a promising possibility for teaching, diagnostic, and research in veterinary pathology.

---

At the end of the 19th century, F. Blum tested the bactericidal properties of a decimal dilution of a commercial solution of formaldehyde (40% w/V) with a small amount of alcohol (usually methanol) used to stabilize the solution and to obtain a 4% w/V solution.^24^ By doing so, he noticed that the skin of his fingers that had been in contact with this solution, became hardened much as with alcohol and curiously analyzed at histology the first formaldehyde-fixed mouse tissue discovering its excellent fixation properties. Since the discovery, this formaldehyde solution has become the most widespread histological fixative all over the world, in both human and veterinary pathology.^24,64^ Commercial formaldehyde is known to have an acidic pH close to 4, but several studies have reported that acidic pH severely damages nucleic acids and that maximum crosslinking and penetration of the fixative occurs in the neutral pH range.^30^ For this reason, in histopathological laboratories, formaldehyde is used in phosphate buffer (1:10) to reach a neutral pH (from 6.8 to 7.2). This solution is called PBF (Phosphate Buffered Formalin) or NBF (Neutral Buffered Formalin). NBF (also named as 10% formalin, which is 3.7% formaldehyde in water with 1% methanol) is the most widely used fixative in pathology owing to its high degree of accuracy and extreme adaptability.^64^ Moreover, the massive worldwide distribution has made NBF also economically extremely competitive.

Despite so many advantages, NBF is an extremely volatile, irritant, and toxic reagent.^41,61,76^ In the early 1980’s, Spengler and Sexton and the World Health Organization identified formaldehyde as one of the priority indoor air pollutants, and in 1987 an indoor guideline level of 0.1 mg/m3 (0.08 ppm) was fixed.^59,61,75^ In 2004, the International Agency for Research on Cancer (IARC) classified formaldehyde as carcinogenic to humans (Group 1).^76^ It is well documented that exposure to NBF can cause sensitization by contact, asthma, irritation of the mucous membranes of the eye and upper respiratory tract.^41^ Other studies have reported the toxic effect of NBF on the central nervous system and the urinary system.^4,33,39,54^ It is also known that NBF vapor exposure increases the risk of developing cancer.^1,16^ Since 2017, the European Commission, with the Regulation (EU) n. 895/2014, has classified formaldehyde as a human carcinogen, category 1b, and germ cell mutagen, category 2, and has prohibited its use.^12,58^ Currently, without a valid alternative on the market, the NBF is used with an exception.

In addition to the toxicity, proteins and nucleic acid analyses from NBF-fixed samples are compromised due to the cross-linking fixation process, with important limitations for a molecular approach on the large number of histological archival samples and the need of parallel sampling for different downstream analyses.^13,47^ To be accepted, an alternative non-carcinogenic fixative would likely be an aldehyde, acting in a chemical reaction similar to formaldehyde, cross-linking proteins and nucleic acids comparably. This should allow histology, histochemistry, immunohistochemistry and molecular procedures to be performed with minor adjustments, so that most of the internationally acquired and accepted diagnostic parameters would not be lost.^12^

Over the years, several alternatives have been proposed to replace NBF but, so far, none of the reactants have shown to be suitable due to problems related to tissue conservation, cell morphology, loss of antigenicity, and high purchase costs.^18,29,46^ To address this antique matter of the use of NBF in pathology, a non-carcinogenic dialdehydic acid-free glyoxal-based fixative (GAF) has been recently proposed.^12^ Over the last decades, several glyoxal-based fixatives have been developed, but a series of unresolved unsatisfactory results (*e.g.*, red blood cell lysis, microcalcification dissolution, unsatisfactory fluorescence in-situ hybridization (FISH), poor nucleic acid preservation) discouraged their use.^20,25,38,69^ GAF is obtained by removing acids from the dialdehyde glyoxal applying a ion-exchange resin, is CE registered, patented, stabilized in a pH 7,1-7,8 in phosphate buffer, and added with phenol red as a pH indicator.^11,12^

Preliminary tests have already been carried out to assess GAF on human tissues.^12,14,34,65^ In these studies, parallel fixation with GAF and NBF was performed on human tissues followed by different standard analytical procedures. The results confirmed the non-inferiority of GAF compared to NBF in all the included protocols for histology, histochemistry, immunohistochemistry, nucleic acid extraction and subsequent molecular biology.

The aim of this multicentric study was to test GAF versus NBF on animal tissues both from biopsies and necropsies, mostly from domestic animal species, to assess standard macroscopic, histological, histochemical, immunohistochemical, ultrastructural, and molecular analysis.

## Materials and methods

### Samples

In this multicentric study, fresh tissues were collected either during diagnostic necropsies submitted mainly within a maximum of 24 hours from the time of death to the BCA Dept. of the University of Padua, or during routine surgeries at the Veterinary Science Dept. of the University of Turin or at the San Marco Veterinary Clinic of Padua (Italy). Double sampling of similar-sized portions was obtained for each tissue either by pathologists or by surgeons and was followed by parallel fixation in a 10x volume NBF (cod. 1617, Kaltek s.r.l, Italy) or GAF (Addax Bioscences, Italy http://addaxbio.com) for 24, 48 or more than 48 hours. Surgical samples were included in the study when multiple biopsies were taken from the same lesion or when the sample was large enough to be divided in half without affecting the diagnosis.

Both NBF and GAF-fixed samples were then trimmed (?), placed in tissue cassettes, processed, and stained with hematoxylin and eosin following the internal standard of each institution (BCA Dept., Padua; Veterinary Science Dept., Turin; San Marco Veterinary Clinic, Padua). No modifications of trimming protocols and processing schedules were applied in order to test GAF performances under routine circumstances.

At the beginning of the study on a small subset of post-mortem samples (5 cases), 2 senior ECVP-certified pathologists and a junior pathologist from the BCA Dept. blindly evaluated the samples without knowing the fixative. The evaluated features were cellular definitions and details, and contrast and brightness of the staining. Since these preliminary results were comparable between the two fixative and among the pathologists (data not shown), with the progression of the study and the involvement of the two additional institutions, the aim became to precisely assess any subtle difference between GAF and NBF. Therefore, at least two pathologists per institution blindly revised the stained slides, evaluating the diagnostic adequacy and annotating any difference between GAF and NBF samples. In case of disagreement between pathologists a final consensus was obtained after joint revision.

Additionally, when required for the diagnosis, variable histochemical and immunohistochemical staining were performed on both GAF and NBF samples according to the standard protocols of each institution.

As per Directive 2010/63/EU of the European Parliament and of the Council of September 22, 2010, regarding the protection of animals used for scientific purposes, the Italian legislature (D. Lgs. n. 26/2014) does not require approval from ethical committees for the use of samples submitted or taken for diagnostic purposes. When submitting the cadavers or before surgery owners signed an informed consent statement to use the samples for research. No additional biopsies were performed at surgery specifically for the study.

### Electron microscope

From one necropsied horse with a suspect of *Herpesvirus* sp. infection, samples from lung and lymph node were submitted for transmission electron microscopy (TEM) after both GAF and NBF fixation and paraffin embedding. The samples were deparaffinized with xylene and hydrated by decreasing scale of ethanol. Subsequently, samples were treated with 2.5% glutaraldehyde solution in 0.1M sodium cacodylate buffer pH 7.4 ON at 4°C. Subsequently they were postfixed with 1% osmium tetroxide in 0.1M sodium cacodylate buffer for 2 hours at 4°. After three water washes, samples were dehydrated in a graded ethanol series and embedded in an epoxy resin (Sigma-Aldrich). Ultrathin sections (60-70 nm) were obtained with an Ultratome Leica Ultracut EM UC7 ultramicrotome, counterstained with uranyl acetate and lead citrate and viewed with a Tecnai G^2^ (FEI) transmission electron microscope operating at 100 kV. Images were captured with a Veleta (Olympus Soft Imaging System) digital camera.

### Nucleic acid analysis

To test nucleic acid preservation in GAF-fixed samples compared to NBF-fixation, DNA was extracted after routine paraffin embedding (2 sections of 20 μm per sample) from 10 canine tumors (DNeasy Blood & Tissue Kit, Cat. No.69504, Qiagen; Veterinary Science Dept. of the University of Turin), 2 canine gastrointestinal biopsies and 5 canine normal tissues (High Pure PCR Template Preparation Kit, Roche Applied Science; San Marco Veterinary Clinic of Padua) according to kit manufacturer instructions. Quantification of extracted DNA was performed by fluorometry (Qubit dsDNA BR assay kit; Qubit 2.0 Fluorometer, Thermo Fischer, CA, USA) and fragmentation analyzed by 1% agarose gel electrophoresis and High Sensitivity Bioanalyzer 2100 assay (Agilent Technologies).

Electrophoresis and gene amplification of canine *TP53* (exon 7) and canine *COX1* were also performed to test DNA fragmentation and efficiency of PCR on subsets of samples. *TP53* was tested on the 5 canine tumoral samples (Veterinary Science Dept. of the University of Turin; primers: sense 3′-TGATAGACTACAGGCCTGCC; antisense 5′-ACAGGAATGGATGGGAAGGA), whereas *COX1* amplification on the 5 normal canine tissues and two canine gastrointestinal biopsies (San Marco Veterinary Clinic of Padua, primers: sense 3′-GGGGCTTTGGAAACTGACTA; antisense 5′-TGGAGGAAGGAGTCAGAAGC). Briefly, PCR was performed using HotStar Taq (Qiagen) at 58◦C (annealing temperature) for 35 cycles, and amplification products were evaluated on agarose gel.

Additionally, due to the suspect of *Herpesvirus sp.* infection in a necropsied horse, DNA was extracted (DNeasy Blood and Tissue Kit/ QIA amp DNA FFPE tissue kit, QIAGEN; BCA Dept. University of Padua) from both GAF and NBF-fixed lung, liver, and lymph-node for viral amplification and sequencing performed at the Istituto Zooprofilattico Sperimentale delle Venezie (Padua, Italy). Briefly, amplification of all samples was performed using a high-fidelity polymerase (Phusion Hot Start II DNA Polymerase, Thermo Scientific) following a consensus primer PCR method for amplification of a region of herpesviral genomes which can be used to detect and partially identify herpesviruses present in tissue samples. (Vandevanter et al., 1996) PCR products were purified by exoSA-IT™ (USB Corp.) and Sanger sequencing was carried out on an automated sequencer (Applied Biosystems). All sequences obtained were aligned with the basic local alignment search tool (BLAST, https://blast.ncbi.nlm.nih.gov/Blast.cgi).

RNA analysis was performed on selected GAF and NBF-fixed canine tumoral samples. RNA was extracted from 3 paraffin 10 um-thick sections per sample. AllPrep DNA/RNA FFPE Kit (Qiagen, Hilden, Germany) was used following the manufacturer’s protocol and the deparaffinization method with xylene, performing also the optional DNAse treatment (Qiagen) for RNA purification. Quality and concentration of obtained RNA was assessed firstly using NanoDrop ND-1000 spectrophotometer (NanoDrop Technologies, Wilmington, USA) (data not shown) followed by the more accurate Qubit Fluorometer (Invitrogen, Carlsbad, CA, USA) and 1% agarose gel electrophoresis.

### RNAScope

Samples of canine mast cell tumor (MCTs) NBF and GAF-fixed paraffin-embedded at the Veterinary Science Dept. of Turin were sent to the Veterinary Medicine and Animal Production Dept., University of Naples, Italy, for RNAScope analysis. From the blocks, 4 μm tissue sections were stained with haematoxylin and eosin (HE) for morphology and with toluidine blue (TB) (code no. T3260, Merck KGaA, Darmstadt, Germany) for the detection of mast cell metachromatic granules. Manual RNAscope assay was performed using BaseScopeTM v2 Assay (cod. # 322350, Bio-Techne, Milan, Italy) according to the manufacturer’s protocol. The RNAScope assay consists of target probes and a signal amplification system composed of a preamplifier, amplifier, and label probe (2,3,4). In the first step, tissues are fixed, and permeabilized to allow the access of the target probe. In the second step, target RNA-specific oligonucleotide probes (conceptualized as a “Z”) are hybridized in pairs (“ZZ”) to multiple RNA targets. In a third step, the detection is carried out by specific binding of oligonucleotide preamplifier molecules linked to several amplifiers containing multiple chromogenic labels. In the last step, signals are detected by developing a chromogen to produce small punctate dots that can provide a quantitative and measurable result. Importantly, the preamplifier cannot bind to a single Z probe (non-paired Z probe) because a Z pair is necessary to bind the preamplifier and generate signals (5). Briefly, tissue sections were baked for 1 h at 60°C, deparaffinized, and treated with Pretreat 1 (Bio-techne, Milan, Italy) for 10 min at room temperature (RT). Target retrieval was performed for 15 min at 100–104°C, followed by protease treatment for 15 min at 40°C. Probes were then hybridized for 2 h at 40°C followed by RNAscope amplification followed by red chromogenic detection. Lastly, the sections were counterstained with hematoxylin and mounted with Bio-Mount (Bio-Techne, Milan, Italy). In this study, the following RNAscope probes were used: *c-KIT* (cod. #512801, Bio-Techne, Milan, Italy) probe encodes for *c-KIT* mRNA that may be detected both in the cytoplasm and nuclei, CI-PPIB (cod. # 437441, Bio-Techne, Milan, Italy) as positive control probe, and dihydrodipi-colinate reductase (dapB), a bacterial gene (cod. #310043, Bio-Techne, Milan, Italy) as negative control probe. PPIB, which encodes for a cyclosporine-binding protein (cyclophilin B), is expressed at a sufficiently low level in most tissues; hence, it is the recommended positive control (6).

### Statistical analyses

Statistical analyses were performed to assess any association or differences among tissue preservation, as indicated asby GAF scoring, and other variables such as time of fixation, size and type of tissue. Statistical analysis was performed using GraphPad Prism 8 software. The Shapiro-Wilk test was used to check normality. Differences between two groups were tested with the two-tailed unpaired Student’s t-test when data were normally distributed or the Mann-Whitney test when data were not normally distributed. Level of significance was set at p < 0.05.

## Results

### Histology, Immunohistochemistry, and TEM

A total of 73 cases obtained either from post-mortem (14 cases, Table 1) or bioptic (Table 2) sampling were included into the study. Samples were collected from 7 different species, with a predominance of canine tissues (60/73). Specifically, 14 necropsies (Table 1) were performed and for each subject double samples from the main organs, with or without lesions, measuring approximately 1.5 cubic cm, were collected for NBF and GAF fixation. Among post-mortem cases, samples from 7 subjects were fixed for <24h, those from 3 subjects were fixed over the weekend (48-72h) and 4 subjects were used to test longer fixation time, for >72h and <5 days. Surgical samples (Table 2) from 59 subjects included variable tissues and lesions and measured 1.5 cubic cm as maximum major size. Among them, 3 groups of samples were randomly created according to fixation times depending on when the surgery was performed and on the planned processing schedule of each lab: <24h (13/59 samples), >24h and ≤48h (20/59 samples), >48h (26/59 samples). Within the latter group of samples, fixation time was for 3 (8 samples), 4 (3 samples), 5 (13 samples), or 6 days (1 sample).

**Table 1.**
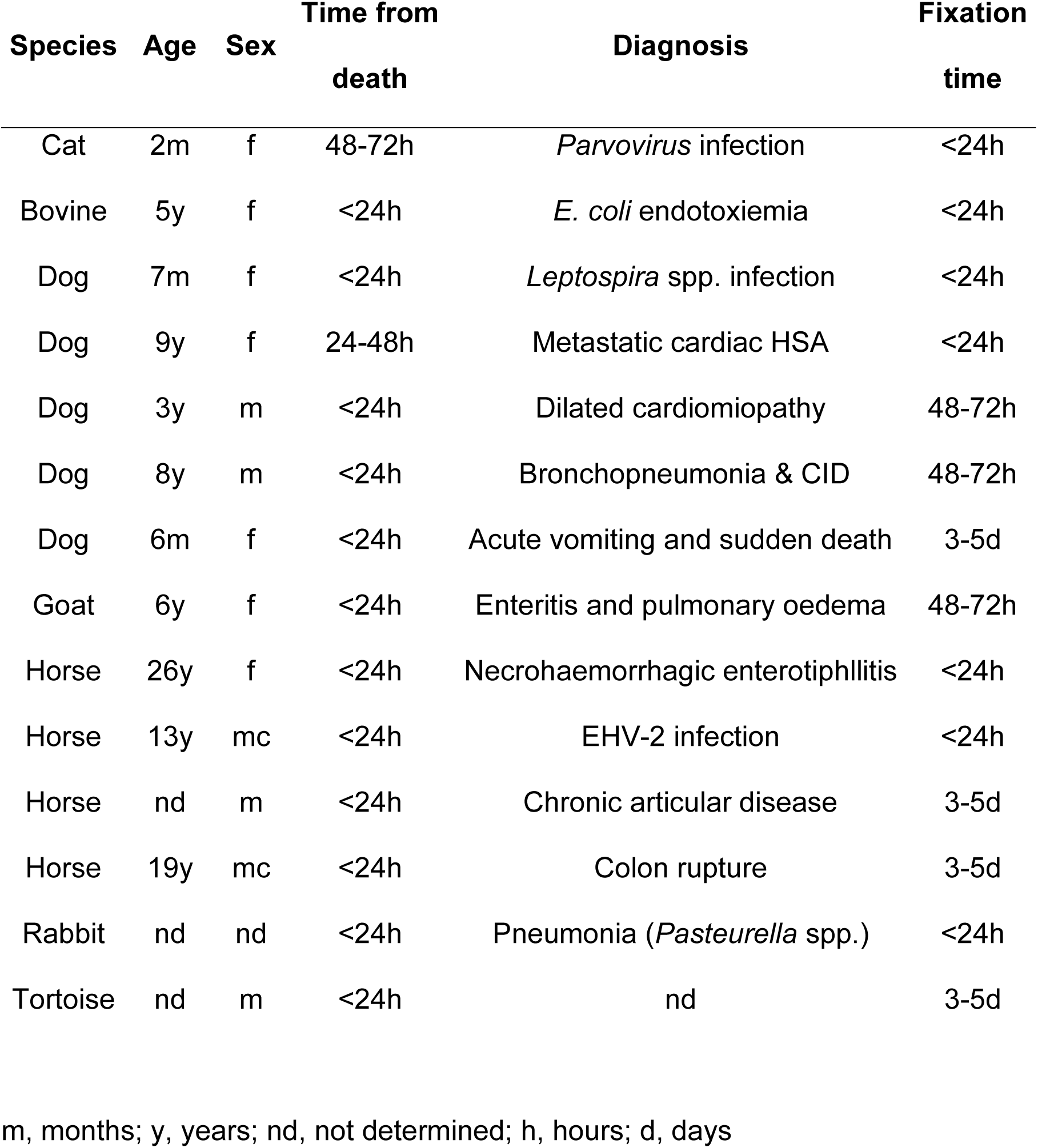
Necropsies included in the study (ordered by species).

**Table 2.**
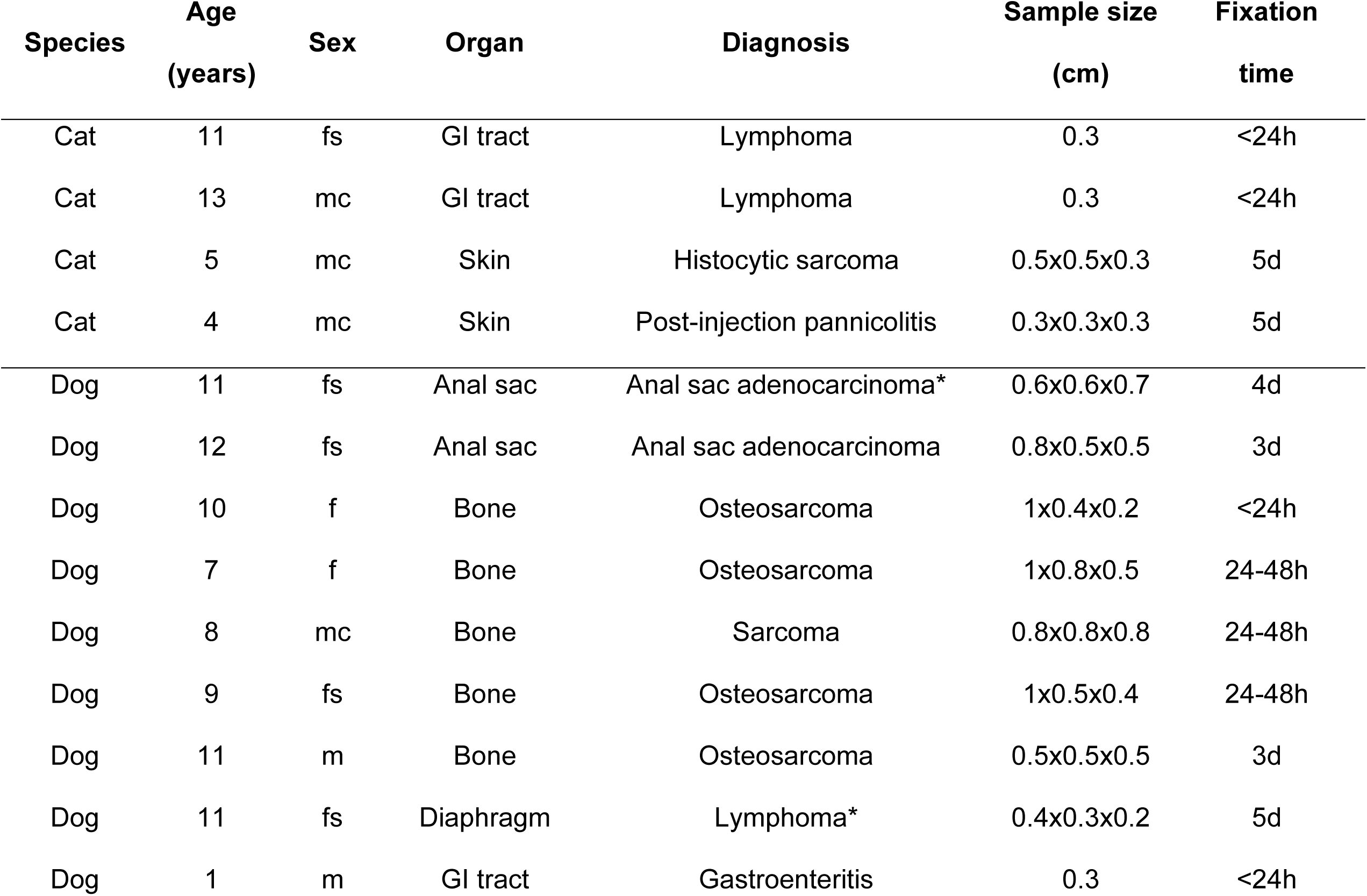
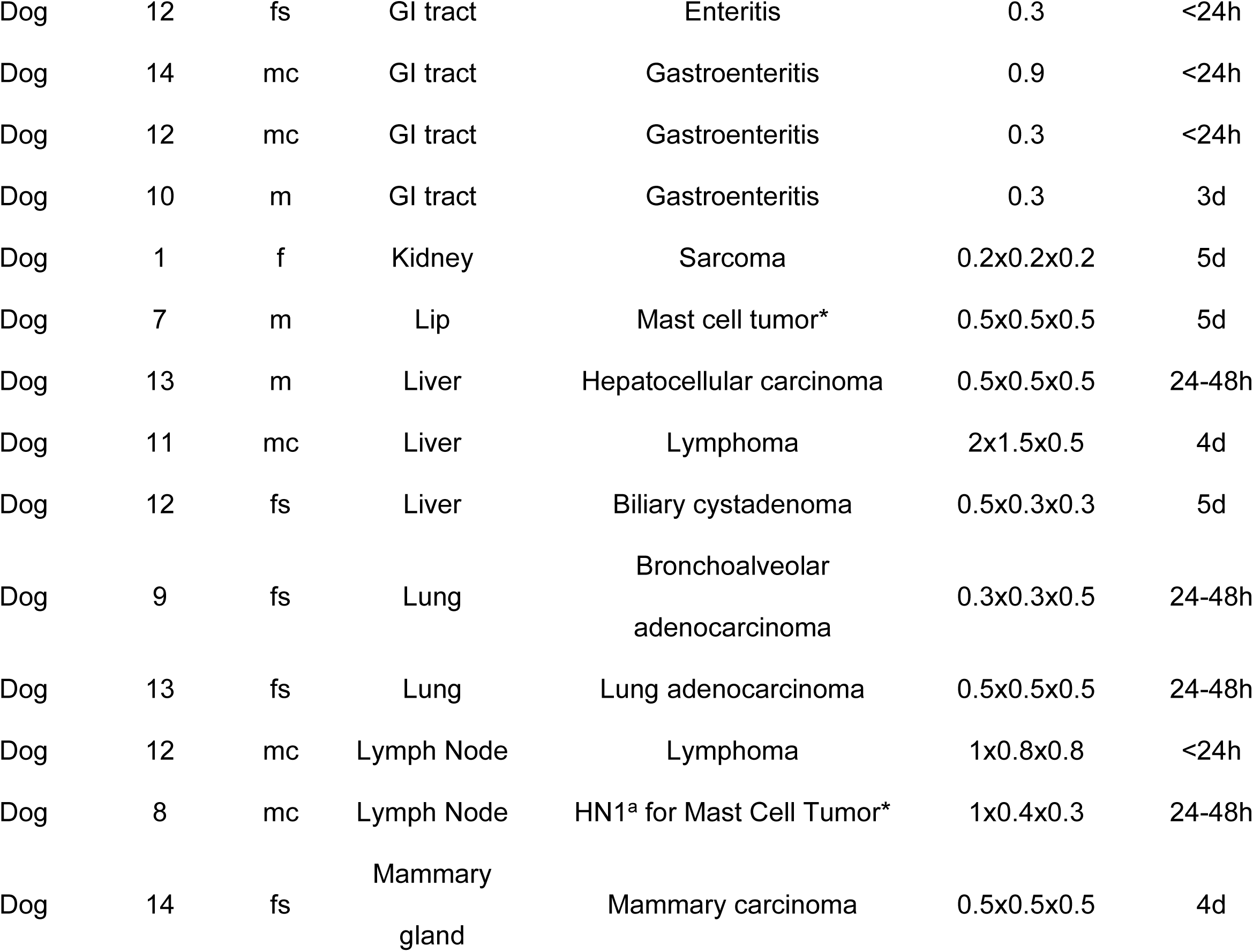
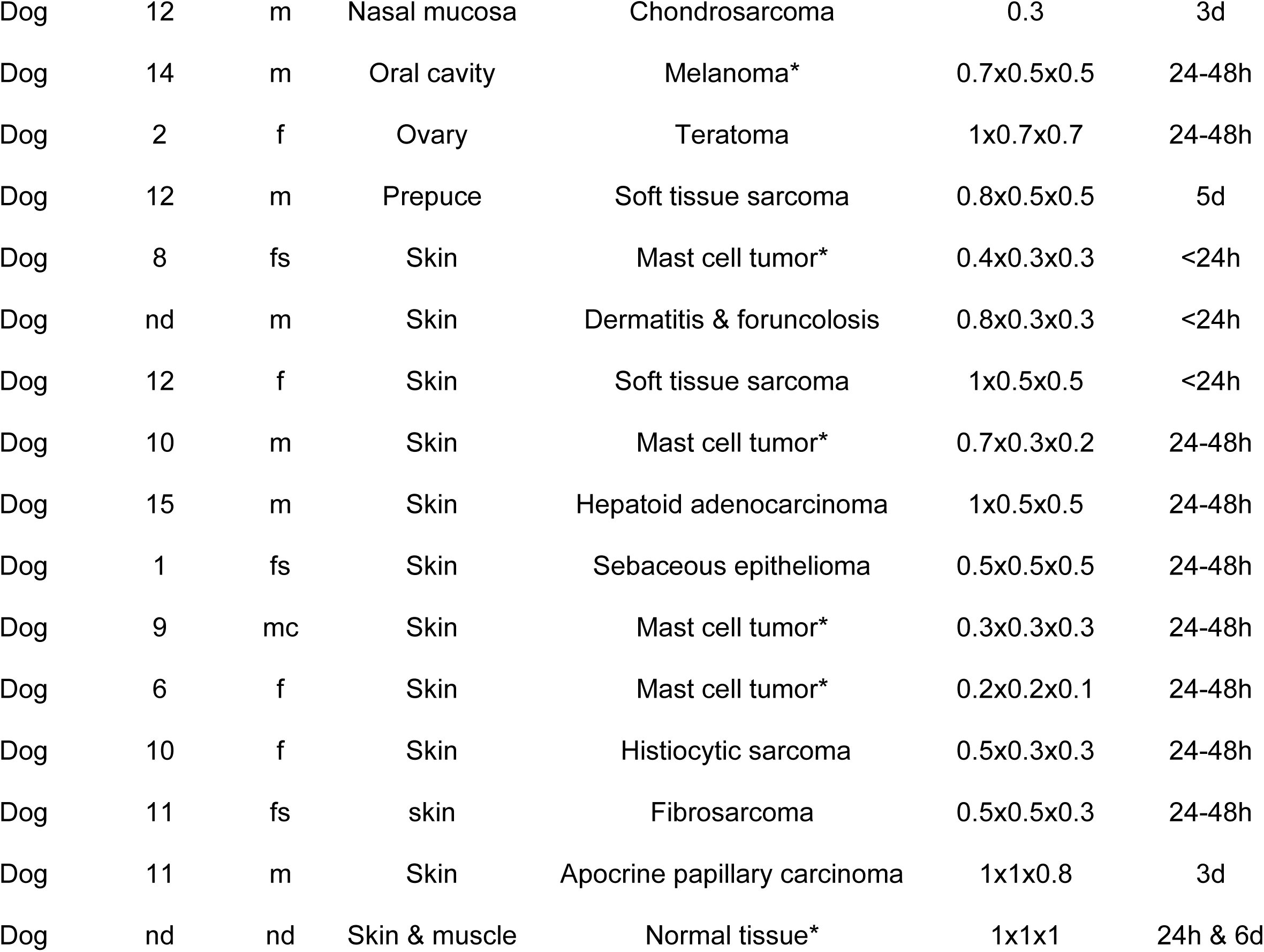
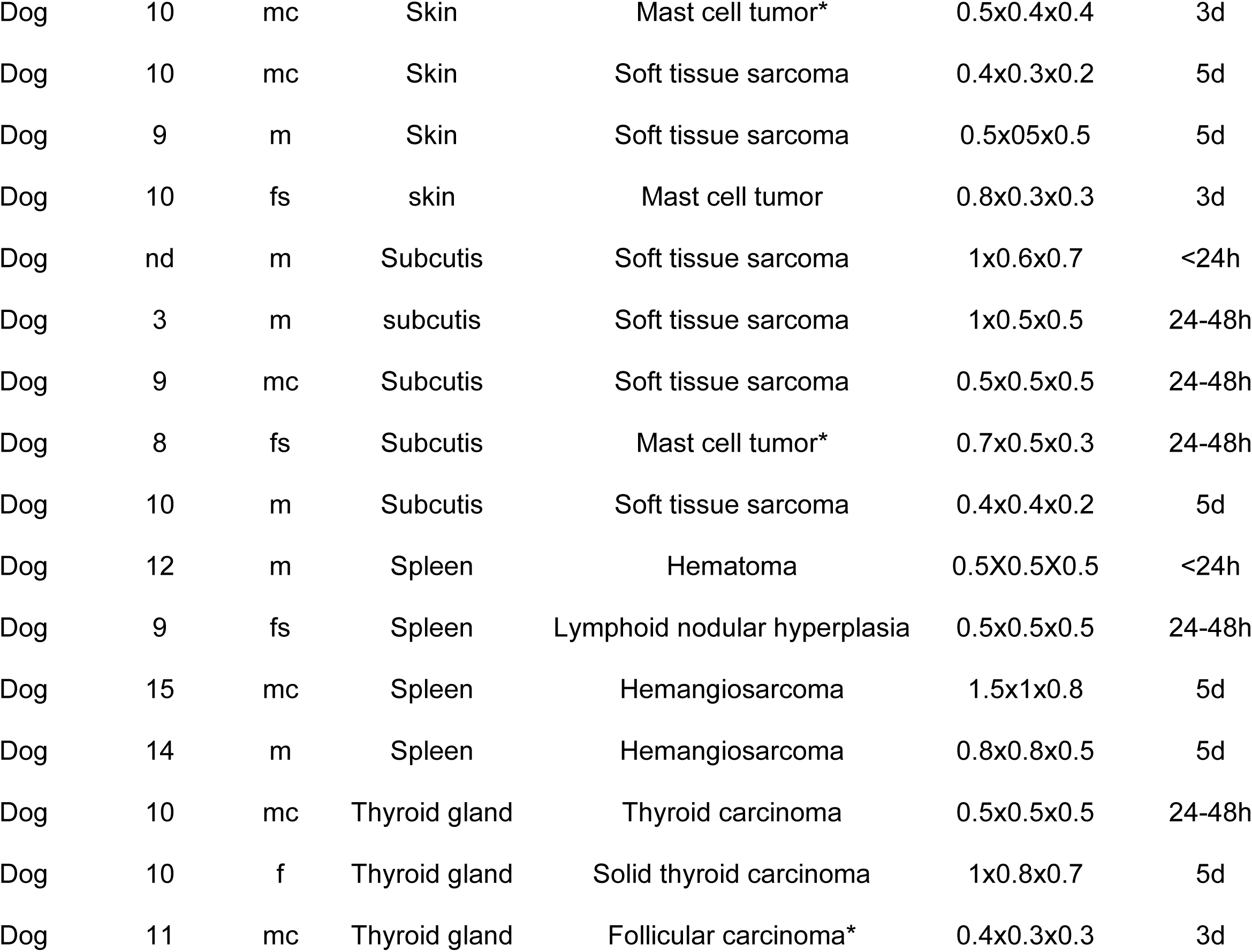
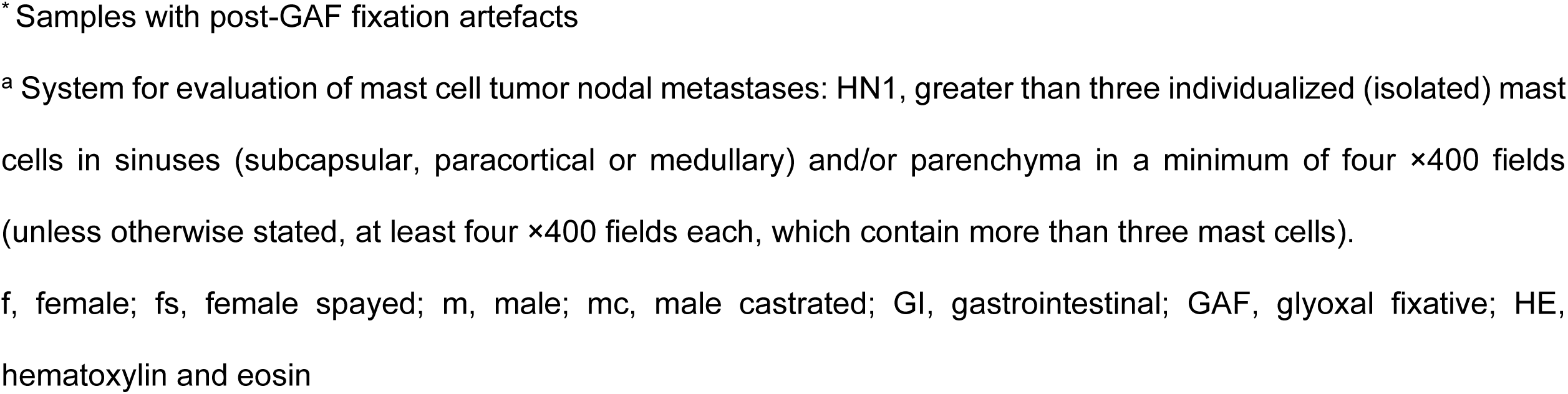
Biopsies included in the study (ordered by species and organ).

| **Species** | **Age**<br>**(years)** | **Sex** | **Organ** | **Diagnosis** | **Sample size**<br>**(cm)** | **Fixation**<br>**time** |
| --- | --- | --- | --- | --- | --- | --- |
| Cat | 11 | fs | GI tract | Lymphoma | 0.3 | <24h |
| Cat | 13 | mc | GI tract | Lymphoma | 0.3 | <24h |
| Cat | 5 | mc | Skin | Histocytic sarcoma | 0.5x0.5x0.3 | 5d |
| Cat | 4 | mc | Skin | Post-injection panniculitis | 0.3x0.3x0.3 | 5d |
| Dog | 11 | fs | Anal sac | Anal sac adenocarcinoma* | 0.6x0.6x0.7 | 4d |
| Dog | 12 | fs | Anal sac | Anal sac adenocarcinoma | 0.8x0.5x0.5 | 3d |
| Dog | 10 | f | Bone | Osteosarcoma | 1x0.4x0.2 | <24h |
| Dog | 7 | f | Bone | Osteosarcoma | 1x0.8x0.5 | 24-48h |
| Dog | 8 | mc | Bone | Sarcoma | 0.8x0.8x0.8 | 24-48h |
| Dog | 9 | fs | Bone | Osteosarcoma | 1x0.5x0.4 | 24-48h |
| Dog | 11 | m | Bone | Osteosarcoma | 0.5x0.5x0.5 | 3d |
| Dog | 11 | fs | Diaphragm | Lymphoma* | 0.4x0.3x0.2 | 5d |
| Dog | 1 | m | GI tract | Gastroenteritis | 0.3 | <24h |
| Dog | 12 | fs | GI tract | Enteritis | 0.3 | <24h |
| Dog | 14 | mc | GI tract | Gastroenteritis | 0.9 | <24h |
| Dog | 12 | mc | GI tract | Gastroenteritis | 0.3 | <24h |
| Dog | 10 | m | GI tract | Gastroenteritis | 0.3 | 3d |
| Dog | 1 | f | Kidney | Sarcoma | 0.2x0.2x0.2 | 5d |
| Dog | 7 | m | Lip | Mast cell tumor* | 0.5x0.5x0.5 | 5d |
| Dog | 13 | m | Liver | Hepatocellular carcinoma | 0.5x0.5x0.5 | 24-48h |
| Dog | 11 | mc | Liver | Lymphoma | 2x1.5x0.5 | 4d |
| Dog | 12 | fs | Liver | Biliary cystadenoma | 0.5x0.3x0.3 | 5d |
| Dog | 9 | fs | Lung | Bronchoalveolar<br>adenocarcinoma | 0.3x0.3x0.5 | 24-48h |
| Dog | 13 | fs | Lung | Lung adenocarcinoma | 0.5x0.5x0.5 | 24-48h |
| Dog | 12 | mc | Lymph Node | Lymphoma | 1x0.8x0.8 | <24h |
| Dog | 8 | mc | Lymph Node | HN1<sup>a</sup> for Mast Cell Tumor* | 1x0.4x0.3 | 24-48h |
| Dog | 14 | fs | Mammary<br>gland | Mammary carcinoma | 0.5x0.5x0.5 | 4d |
| Dog | 12 | m | Nasal mucosa | Chondrosarcoma | 0.3 | 3d |
| Dog | 14 | m | Oral cavity | Melanoma* | 0.7x0.5x0.5 | 24-48h |
| Dog | 2 | f | Ovary | Teratoma | 1x0.7x0.7 | 24-48h |
| Dog | 12 | m | Prepuce | Soft tissue sarcoma | 0.8x0.5x0.5 | 5d |
| Dog | 8 | fs | Skin | Mast cell tumor* | 0.4x0.3x0.3 | <24h |
| Dog | nd | m | Skin | Dermatitis & forunculosis | 0.8x0.3x0.3 | <24h |
| Dog | 12 | f | Skin | Soft tissue sarcoma | 1x0.5x0.5 | <24h |
| Dog | 10 | m | Skin | Mast cell tumor* | 0.7x0.3x0.2 | 24-48h |
| Dog | 15 | m | Skin | Hepatoid adenocarcinoma | 1x0.5x0.5 | 24-48h |
| Dog | 1 | fs | Skin | Sebaceous epithelioma | 0.5x0.5x0.5 | 24-48h |
| Dog | 9 | mc | Skin | Mast cell tumor* | 0.3x0.3x0.3 | 24-48h |
| Dog | 6 | f | Skin | Mast cell tumor* | 0.2x0.2x0.1 | 24-48h |
| Dog | 10 | f | Skin | Histiocytic sarcoma | 0.5x0.3x0.3 | 24-48h |
| Dog | 11 | fs | skin | Fibrosarcoma | 0.5x0.5x0.3 | 24-48h |
| Dog | 11 | m | Skin | Apocrine papillary carcinoma | 1x1x0.8 | 3d |
| Dog | nd | nd | Skin & muscle | Normal tissue* | 1x1x1 | 24h & 6d |
| Dog | 10 | mc | Skin | Mast cell tumor* | 0.5x0.4x0.4 | 3d |
| Dog | 10 | mc | Skin | Soft tissue sarcoma | 0.4x0.3x0.2 | 5d |
| Dog | 9 | m | Skin | Soft tissue sarcoma | 0.5x0.5x0.5 | 5d |
| Dog | 10 | fs | skin | Mast cell tumor | 0.8x0.3x0.3 | 3d |
| Dog | nd | m | Subcutis | Soft tissue sarcoma | 1x0.6x0.7 | <24h |
| Dog | 3 | m | subcutis | Soft tissue sarcoma | 1x0.5x0.5 | 24-48h |
| Dog | 9 | mc | Subcutis | Soft tissue sarcoma | 0.5x0.5x0.5 | 24-48h |
| Dog | 8 | fs | Subcutis | Mast cell tumor* | 0.7x0.5x0.3 | 24-48h |
| Dog | 10 | m | Subcutis | Soft tissue sarcoma | 0.4x0.4x0.2 | 5d |
| Dog | 12 | m | Spleen | Hematoma | 0.5X0.5X0.5 | <24h |
| Dog | 9 | fs | Spleen | Lymphoid nodular hyperplasia | 0.5x0.5x0.5 | 24-48h |
| Dog | 15 | mc | Spleen | Hemangiosarcoma | 1.5x1x0.8 | 5d |
| Dog | 14 | m | Spleen | Hemangiosarcoma | 0.8x0.8x0.5 | 5d |
| Dog | 10 | mc | Thyroid gland | Thyroid carcinoma | 0.5x0.5x0.5 | 24-48h |
| Dog | 10 | f | Thyroid gland | Solid thyroid carcinoma | 1x0.8x0.7 | 5d |
| Dog | 11 | mc | Thyroid gland | Follicular carcinoma* | 0.4x0.3x0.3 | 3d |

\* Samples with post-GAF fixation artefacts
<sup>a</sup> System for evaluation of mast cell tumor nodal metastases: HN1, greater than three individualized (isolated) mast cells in sinuses (subcapsular, paracortical or medullary) and/or parenchyma in a minimum of four ×400 fields (unless otherwise stated, at least four ×400 fields each, which contain more than three mast cells).
f, female; fs, female spayed; m, male; mc, male castrated; GI, gastrointestinal; GAF, glyoxal fixative; HE, hematoxylin and eosin

One common result to all samples was that GAF-fixed samples did not show significant gross color or consistency change, as in NBF-fixed samples. Every type of tissue maintained the original consistency, without hardening, and preserved original colors (Fig. 1).

**Figure 1.**
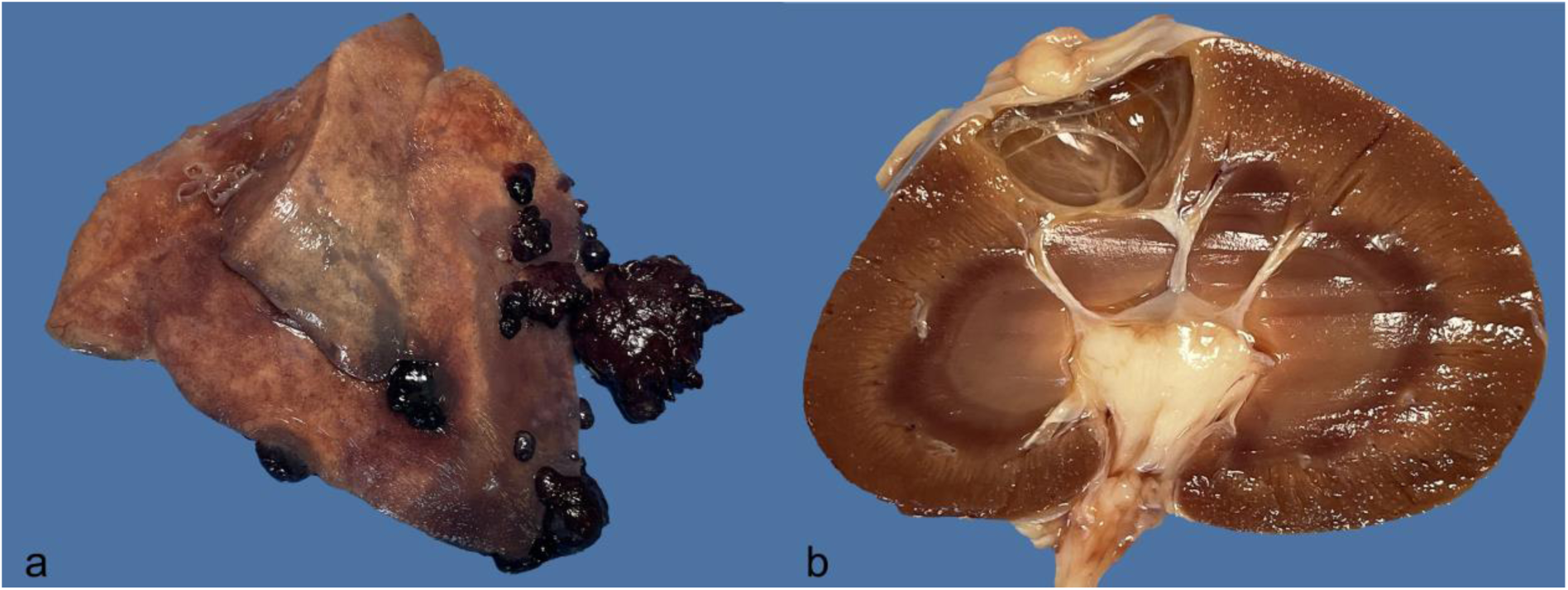
Visceral organs fixed in glyoxal-based fixative (GAF) for 15 days. a) Metastatic hemangiosarcoma in the lung, dog. b) Renal cortical cyst, cat.

Generally, on histology GAF and NBF fixed samples were comparable and considered of adequate quality for histopathological diagnoses by all pathologists from the 3 institutions (Fig. 2). One histological difference was the evidence of erythrolysis in some GAF fixed samples particularly in highly congested tissues (Fig. 2d). Additional differences were observed in skeletal and cardiac muscles that were mainly obtained from necropsies and manifested less tissue cracking and splitting with GAF fixation than with NBF (Fig. 2f). Also, all mast cell tumors (8 primary tumors and 1 lymph node metastasis, the latter assessed according to Weishaar and co-authors, 2014 showed a diffuse loss of cytoplasmic staining in mast cells when fixed with GAF (Fig. 3b).^74^ In addition to mast cell tumors, 5 canine surgical samples including an anal sac adenocarcinoma, a lymphoma, an oral melanoma, a cutaneous sample with no lesion, and a thyroid follicular carcinoma, had some artifactual changes compared to NBF, but diagnoses were not impaired. In these cases, the major artifact was the cell shrinkage and detachment from the embedding matrix or from the basement membrane (Table 2). No significant association or differences were detected between tissue preservation (GAF scoring) and time of fixation, type and size of tissue, or other subject/sample related features (*e.g.,* species, age, sex, diagnosis).

**Figure 2.**
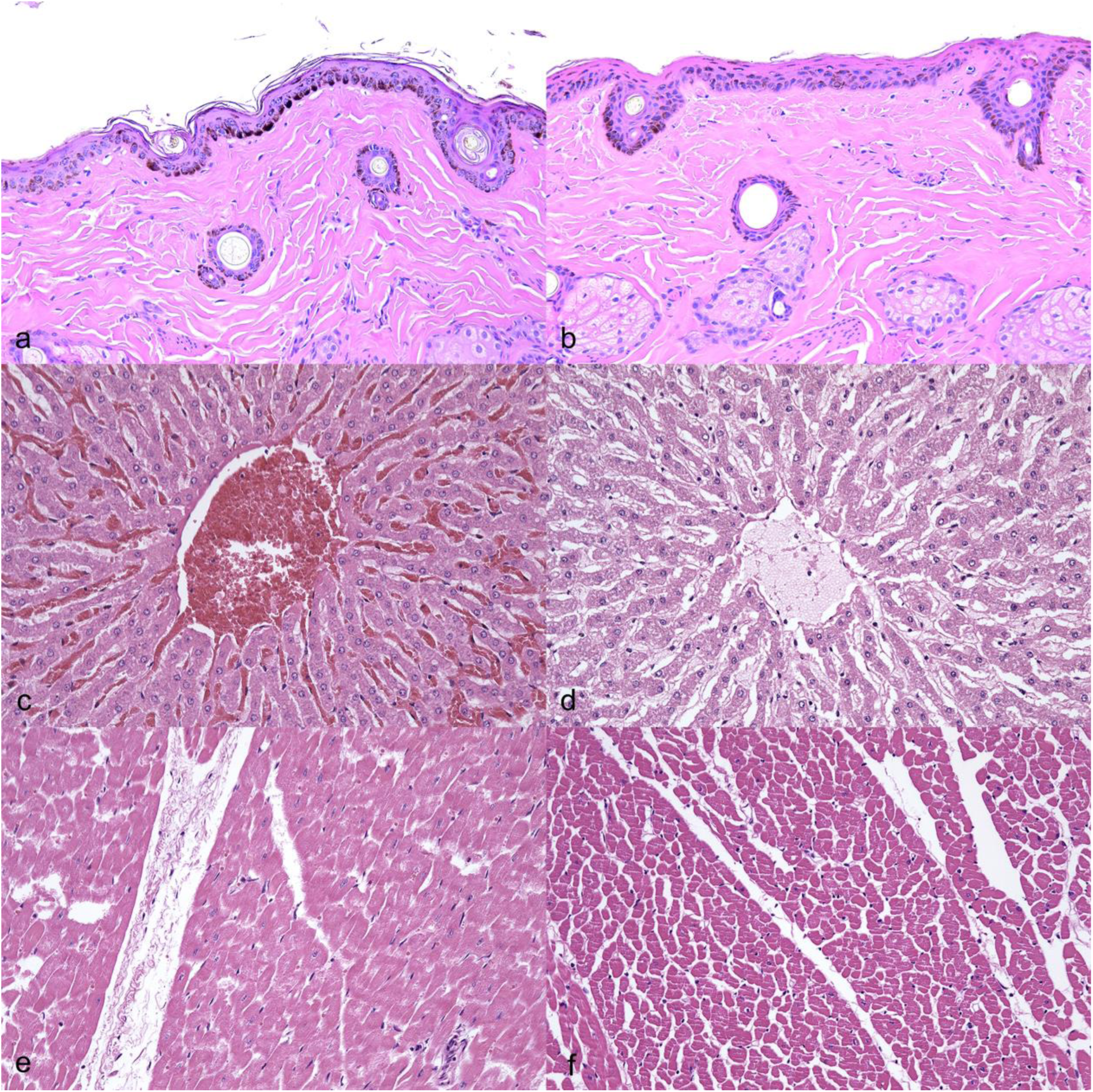
Neutral buffered formaldehyde (NBF) (a, c, e, g) versus glyoxal-based fixative (GAF) (b, d, f, h) fixation. a) Skin, NBF fixation, dog, HE. b) Skin, GAF fixation, dog, HE. Note very similar morphological features when compared to formalin. c) Liver, NBF fixation, dog, HE. d) Liver, GAF fixation, dog, HE. Note some degree of erythrolysis. e) Skeletal muscle, NBF fixation, dog, HE. f) Skeletal muscle, GAF fixation, dog, HE. Note less tissue cracking when compared to formalin.

**Figure 3.**
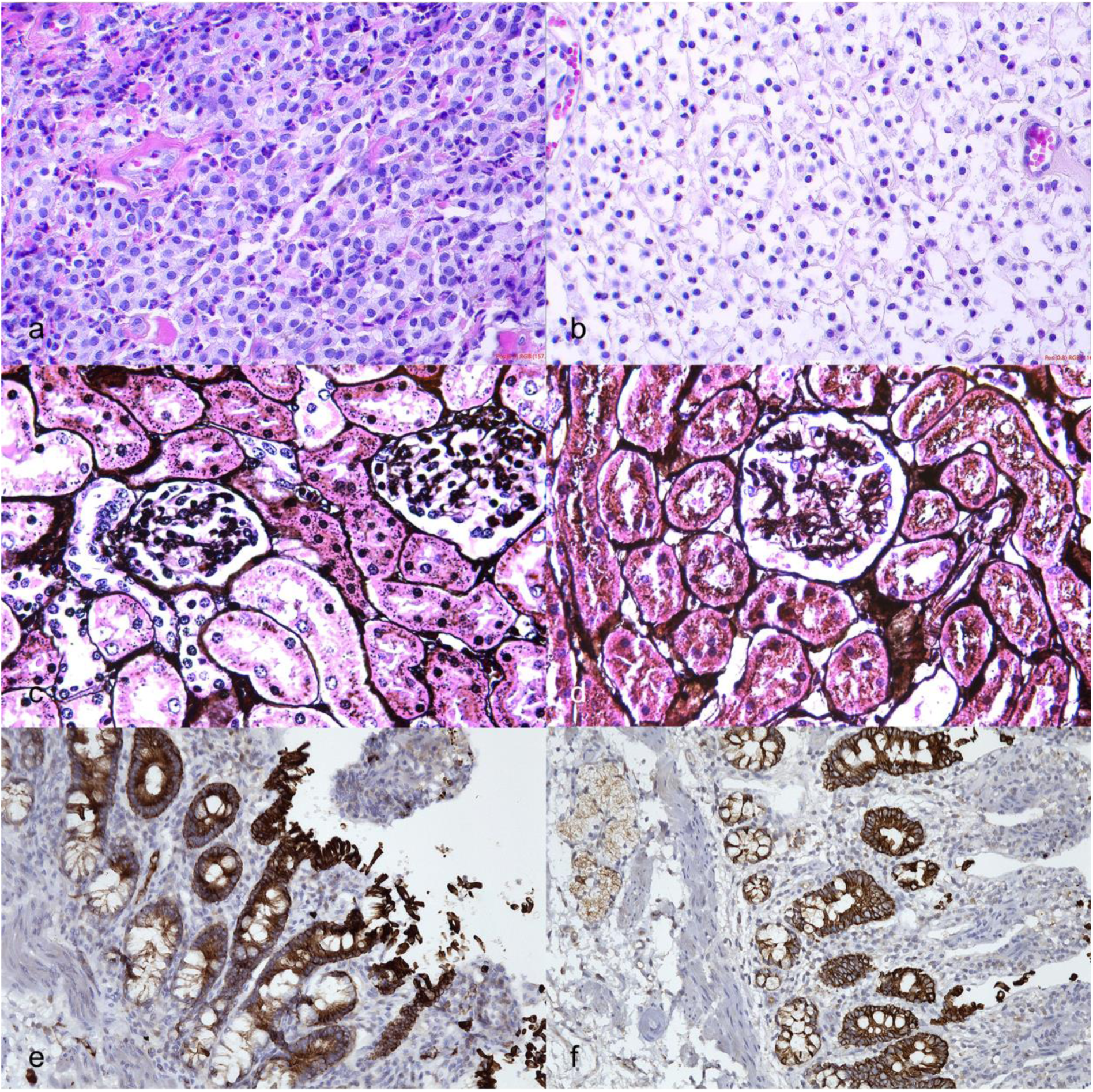
Neutral buffered formaldehyde (NBF) (a, c, e) versus glyoxal-based fixative (GAF) (b, d, f) fixation. a) Cutaneous mast cell tumor, NBF fixation, dog, HE. b) Cutaneous mast cell tumor, GAF fixation, dog, HE. Note that mast cells lose the basophilic granular cytoplasm. c) Kidney, NBF fixation, dog, Periodic Schiff-Methenamine Silver (PASM). d) Kidney, GAF fixation, dog, PASM. Note very similar morphological features when compared to formalin. e) intestine, NBF fixation, dog, pancytokeratins immunohistochemistry, DAB. f) Intestine, GAF fixation, dog, pancytokeratins immunohistochemistry, DAB. Note very similar specific and sensitivity when compared to NBF.

A total of 18 tissues underwent additional histochemical staining including silver stain, Giemsa, Gram, Grocot, Masson Trichrome, Periodic Acid-Schiff (PAS), Periodic Schiff-Methenamine Silver (PASM), Toloudine Blue, and Zheil Nielsen, performed according to routine protocols. Results were always comparable between NBF and GAF fixation (Figs. 3c and 3d). Immunohistochemistry was carried out on a total of 44 samples for the following antigens: alpha smooth muscle actin, calcitonin, CD3, CD20, CD31, c-Kit, cytokeratins, IBA1, Ki-67, p63, thyroglobulin, vimentin, von Willebrand Factor. Most routinely applied, both automized (Benchmark XT, Ventana Medical System) and manual protocols (not presented, available from the Institutions), gave optimal overlapping results among GAF and NBF fixed samples (Figs. 3e and 3f) except for three antigens (Ki-67, IBA1 and c-Kit) that performed less optimally in GAF fixed samples, with less intense staining of a lower number of cells. The protocol for Ki-67 was slightly adjusted according to GAF manufacturer available protocol (https://addaxbio.com/ihc-protocols/) and an improvement of results was obtained (data not shown).

At TEM, performed on equine paraffin-embedded samples, GAF fixation seemed to better preserve cell membranes and integrity with also less chromatin and organelles dissolution (Figs. 4a and 4b). No viral particles were detected.

**Figure 4.**
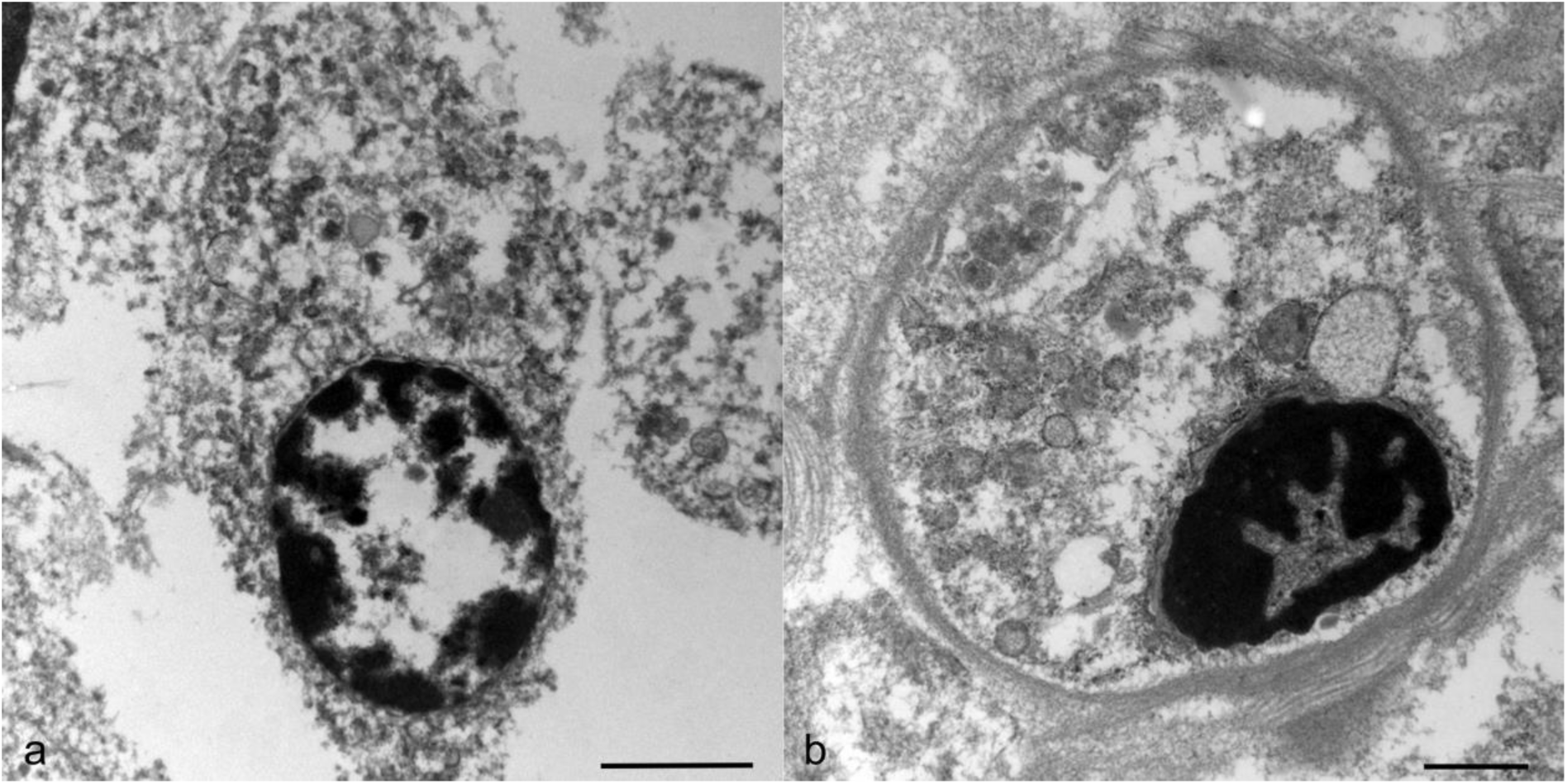
Neutral buffered formaldehyde (NBF) versus acid-free glyoxal-based fixative (GAF) at transmission electron microscopy (TEM). a) Lung, NBF-fixation, horse, transmission electron microscopy (TEM). b) Lung, GAF-fixation, horse, transmission electron microscopy (TEM). Note better preservation of cell membranes and organelles when compared to NBF.

### Nucleic acid analyses and RNAScope

Regarding DNA analyses, most of the samples showed a higher DNA yield from GAF-fixed tissues (P<0.05) except for one MCT (Table 3). Both tested MCTs showed a lower DNA yield compared to the other samples. When excluding MCTs from the statistical analyses the difference of DNA yield between GAF and NBF fixation raised to P ≤ 0.01 (Fig. 5a). DNA fragmentation did not show significant difference between the two fixatives showing for both of them >50% of fragments superior to 10000bp (data not shown).

**Table 3.** DNA and RNA extraction from paraffin-embedded glyoxal (GAF)-fixed and formalin (PBF)-fixed canine samples.

|  | DNA (ng/ul) |  | RNA (ng/ul) |  |
| --- | --- | --- | --- | --- |
|  | GAF | PBF | GAF | PBF |
| Oral melanoma | 77.6 | 41.8 | 1136.0 | 452.0 |
| Hepatocellular k | 8.8 | 35.6 | 411.0 | 187.8 |
| Soft tissue sarcoma | 89.6 | 41.2 | 590.8 | 144.0 |
| Soft tissue sarcoma | 87.3 | 61.1 | 1008.0 | 620.0 |
| Teratoma | 150.6 | 63.1 | 777.0 | 600.0 |
| Anal sac adenok | 241.2 | 206.4 | 141.0 | 74.5 |
| Thyroid k | 59.5 | 14.5 | 98.6 | 20.8 |
| Aprocrine k | 73 | 3,05 |  |  |
| Lip mct | 3,24 | 1,67 |  |  |
| Cutaneous mct | 1,19 | 2,89 |  |  |
| Liver | 110 | 50 |  |  |
| Kidney | 740 | 160 |  |  |
| Heart | 410 | 100 |  |  |
| Lung | 990 | 0 |  |  |
| Intestine | 1020 | 120 |  |  |
| GI biopsy | 810 | 90 |  |  |
| GI biopsy | 590 | 40 |  |  |

K, carcinoma; mct, mast cell tumor; GI, gastrointestinal

**Figure 5.**
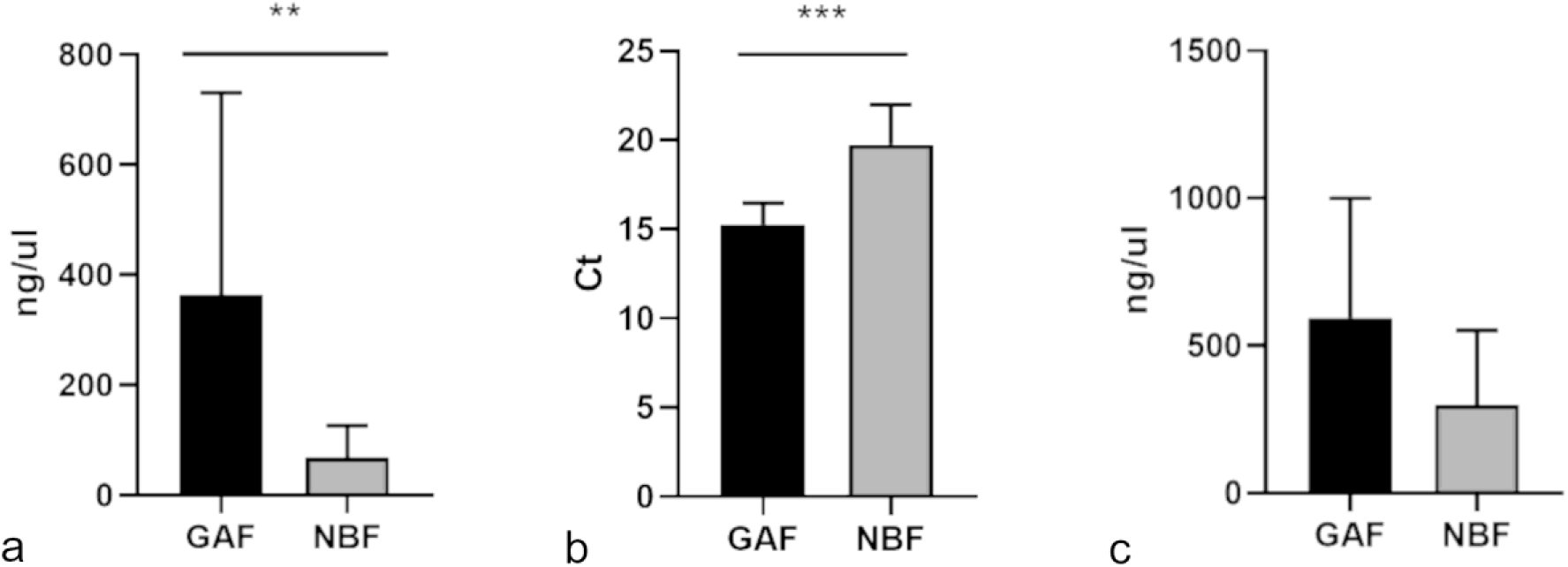
a) DNA extraction showed a higher yield from acid-free glyoxal-based fixative (GAF) samples when compared to neutral buffered formaldehyde (NBF)-fixed samples. b) COX1 gene amplification was performed with significantly lower Ct from GAF-fixed samples then from NBF samples. c) RNA extraction from a subset of samples was higher for GAF-fixed samples then from NBF-fixed samples.

Amplification of *TP53* (exon 7) carried out on 5 canine tumor samples, was successful only from GAF fixed tissues (3/5, excluding 2 MTCs). Further, Ct numbers for amplification of *COX1* from 2 gastrointestinal biopsies and 5 normal canine tissues was significantly lower from GAF fixed samples than NBF (Fig. 5b) again showing a better preservation and higher quantity of DNA after GAF fixation.

Biomolecular evidence of *Equid gammaherpesvirus* 2 (EHV-2) infection was obtained from both GAF and NBF-fixed lung and lymph node and from NBF-fixed liver. From NBF-fixed lung and liver, a 166bp fragment was isolated, presenting 100% homology with the viral sequence Equid herpesvirus 2 strain 86 (GenBank accession number HQ247791). A 166 bp and a 111 bp fragment were identified from GAF preserved lung and lymph node, respectively, with 99% and 97% homology with the viral sequence of the same Equid herpesvirus 2 strain 86 (GenBank accession number HQ247791).

RNA extraction performed from 7 canine tumors also showed a higher yield from all GAF-fixed samples when compared to NBF-fixed samples (Table 3 and Fig. 5c). RNA fragmentation as observed by agarose gel electrophoresis did not show significant difference between the two fixatives and all samples showed most of the fragments having a length superior to 200bp (data not shown).

Eight canine MTCs were submitted for RNAscope. Before processing they were stained with Toluidine blue and metachromasia of cytoplasmic granules was confirmed (Fig. 6a). At RNAScope c-Kit mRNA was positively detected in all GAF samples (Fig. 6b) as well as the internal control CI-PPIB (Fig. 6c), whereas all samples were negative for the bacterial dapB (Fig. 6d).

**Figure 6.**
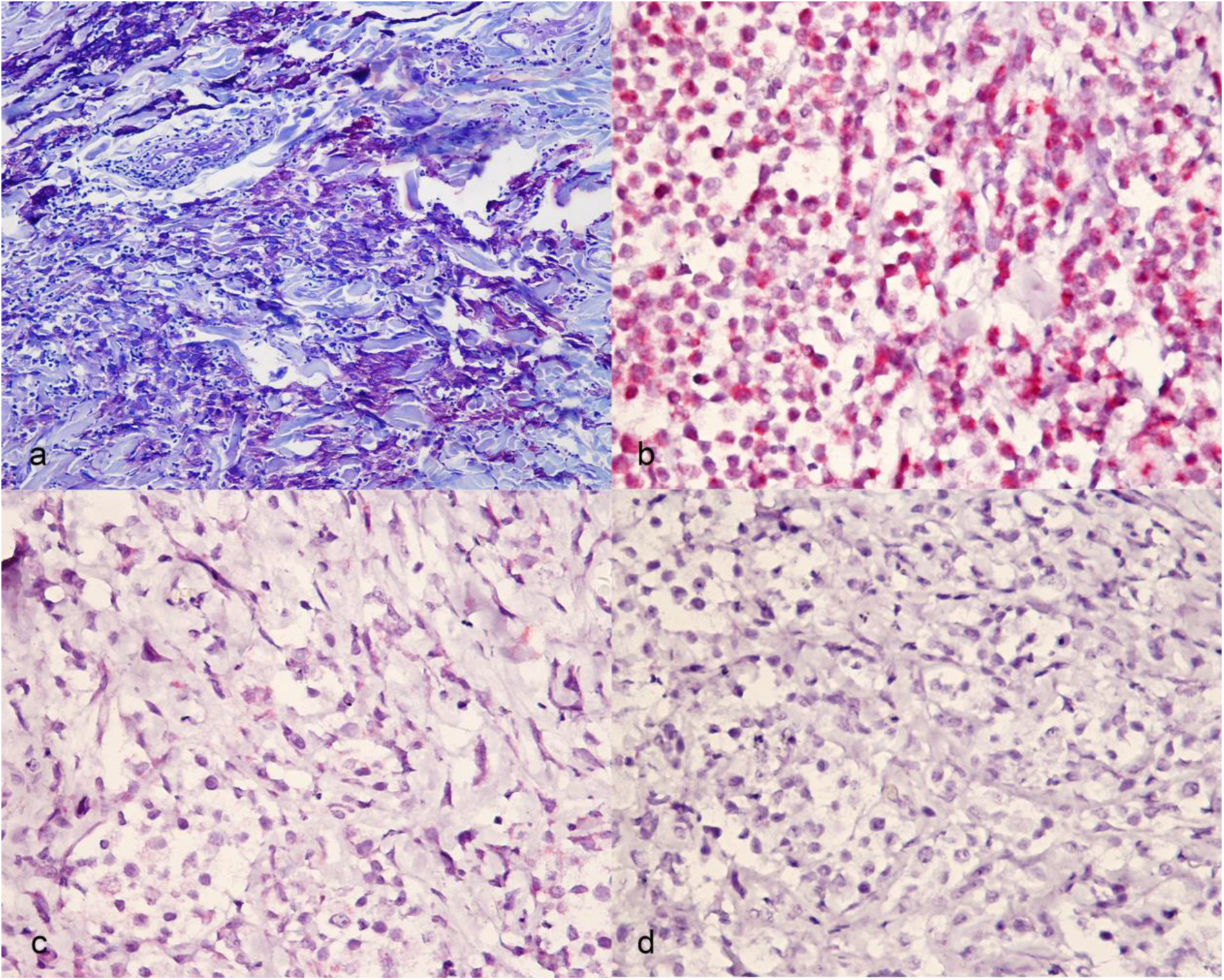
Canine mast cell tumor fixed in acid-free glyoxal-based fixative (GAF) fixed samples. a) Toluidine blue stained metacromatic, purple, intracytoplasmic granules in mast cells. b) c-KIT, c) peptidylprolyl isomerase B d) dihydrodipi-colinate reductase (dapB) expression in mast cells by RNAScope.

## Discussion

In this study the newly-patented, non-toxic, acid-free glyoxal-based fixative (GAF) was tested on animal tissues in comparison with neutral buffered formaldehyde (NBF) for morphology, histochemistry, immunohistochemistry, and nucleic acid analyses. NBF is massively world widely used both in human and veterinary pathology because of its great ability to fix tissues for histology. However, several problems arise with NBF and many strategies have been sought to overcome unwanted NBF effects.^13,55^ Tissue shrinkage and distortion have been widely reported after NBF fixation as well as texture (tissue “hardening”) and color changes are well known by experience from pathologists working with NBF.^2,37,57,77^ Particularly, shrinkage and distortion can affect the evaluation of tissue periphery which is relevant for example when evaluating tumor size or margins or when studying skeletal muscle changes.^2,37,57^ In addition, color and texture changes can affect post-fixation gross examination at trimming or at teaching lab demonstration of non-fresh organ lesions, warranting the use of alternative fixatives known for not altering much texture and colors, such as Klotz.^32^ However, unsatisfactory results and safety issues, make this issue still unsolved. In our study, we detected impressive tissue texture and color preservation with GAF fixation which allowed us to use necropsy organs at teaching labs several days after fixation, also without the need of safety protections at examination (*i.e.,* chemical hood). The maintenance of original texture of GAF-fixed tissues was also noted at trimming and despite changing the perception when slicing the tissue for histological processing, no difficulties or impediments in the procedure were detected. Detailed studies on long term GAF fixation (*e.g.* months, maybe years), trimming of specific organs for which texture is relevant (*i.e.* brain), and precise tissue shrinkage comparison between GAF and NBF should certainly be performed to better assess GAF performances and its applicability for example in oncological studies.

Regarding histology, histochemistry, immunohistochemistry, and molecular analyses our results were similar to what previously reported for human tissues.^12^ All GAF-fixed samples analyzed in this study were considered adequate for the diagnostic process. Some histological limitations in GAF-fixed samples included a certain degree of lysis of erythrocytes in highly congested tissues, the loss of staining of cytoplasmic granules in mast cells, and the occasional shrinkage of stroma/basement membrane. Similar criticisms have been already reported when using commercially available glyoxal as fixative.^12,20,69,72^ Detrimental effects of commercially available glyoxal were, however, much higher than GAF with additional loss of eosinophilia, microcalcification dissolution, and diffuse cellular pallor, giving unsatisfactory results when compared to NBF.^20,42,69,72^ These destructive effects were likely linked to the acidity of the fixative leading to the tendency of the dialdehyde to oxidize and dropping the pH down to 4. Accumulated glyoxylic and other acids hydrolyze molecules disrupting the structure of the tissue.^78^ In order to avoid these damages the dialdehyde glyoxal was removed from acids with an ion-exchange resin creating the neutral GAF, that in the previous and this study gave much more satisfactory results than acidic glyoxal based fixatives.^10,12,52,66,69^ For this reason, an improved GAF pH stabilization is ongoing to further optimize its performances and eliminate the residual unwanted effects. With particular regards to mast cells, they are well known for their acidic cytoplasmic components and their staining is therefore affected by the pH of the applied histochemical method.^44,60^ Additionally, some authors already reported that non-aldehyde fixatives work better for mast cells.^28^ A specific histochemical study could be performed to test variation of staining in mast cell tumors after GAF-fixation. Considering the skeletal and cardiac muscles, in this study we observed less artifacts with GAF then with NBF. Further studies should be carried out to test specific enzymatic staining on GAF-fixed muscular tissues.

Even if performed only on a couple of samples, TEM showed a better preservation of cellular structures with GAF then with NBF. Literature reports based on other glyoxal-based fixatives showed variable results at electron microscopy, indicating that GAF adjustments might have improved the fixation process causing less cross-linking and tissue disruption.^27,56,68^

Immunohistochemistry on GAF-fixed samples also gave results comparable to NBF, with the only exception of three antigens that were less detected after GAF-fixation. These results were similar to what already reported for GAF used with human tissues and superior to the suboptimal results obtained with other glyoxal-based fixatives.^12,19,42,69^ The protocol to detect the nuclear antigen Ki-67 could be easily optimized with minimal modifications (*i.e.* increased antigen retrieval) as already suggested by the manufacturer’s technical sheets, whereas c-Kit and IBA-1 protocols still need adjustments. These two markers, as many others in veterinary pathology, are analyzed applying non-specific anti-human antibodies which make specificity a struggling and very breakable issue. Generally speaking, IHC sensitivity and specificity can be very inconsistent and unpredictable due to pre-analytical, analytical, and post-analytical issues so that standardization is still a matter of concern both in human and veterinary pathology.^3,15,23,40,43,49–51,63,67^ Therefore, it was a positive surprise, that most of the markers were identically detected in GAF and in NBF-fixed samples with unchanged protocols; the minimal required adjustments would be similar to optimization needed when changing an antibody or a specific tissue/species.

With regard to nucleic acid preservation and molecular biology analyses, GAF showed to be a very promising fixative, allowing higher DNA and RNA yields from tissues, a more efficient amplification of tested genes, adequate viral DNA sequencing, and very good results at RNAscope. In addition to this, next generation sequencing and fluorescent in situ hybridization on human samples were also proven to be as equal or better on GAF-fixed samples when compared to NBF.^12,56^

Tissue preservation by NBF is known to limit various methods of genomic analysis and molecular biomarker tests so that parallel sampling is often needed when both morphology and genetic analyses are required on a specific sample.^35,45,73^ NBF is known to affect DNA and RNA integrity by different mechanisms, among which cross-linking of proteins with nucleic acids has been addressed as the major responsible.^62^ Recently, some authors proposed alcoholic fixation as a safer alternative to NBF for morphology and molecular analysis of animal tissues.^53^ Alcoholic fixation is advantageous over formalin fixation because of faster fixation and safer workplace environment also revealing higher total genomic DNA and RNA yields.^53^ However, the protein denaturing process of alcohol-based/non-cross-linking fixatives will never replace the cross-linking aldehyde fixatives such as NBF optimally preserving tissue structure and maintaining both secondary and tertiary structures of proteins.^6,21,31^ Glyoxal-based fixatives reduce the cross-linking of proteins particularly to RNA and guarantee longer DNA fragments also over time.^6,12,14^ We detected similar capacity of viral DNA sequencing when comparing GAF to NBF-fixed samples, whereas the yields of nucleic acids was generally higher from GAF, despite variable among samples, therefore allowing better gene amplification. As a dialdehyde-based non-acid fixative, GAF appears therefore matching both the best structural preservation methodology and a higher nucleic acid maintenance compared to NBF. DNA/RNA variable yields and integrity among samples are reasonably due to different tissue components and features. In this study we randomly included diagnostic samples without assessing features such necrosis or relative amount of extracellular matrix versus cellular components. Further analyses could more carefully assess how pre-analytical features can impact nucleic acid preservation in different fixatives. Lastly, we also detected a positive signal in GAF fixed samples with RNAScope on mast cell tumors. We chose to include mast cell tumors as a preliminary test to evaluate RNA preservation since these samples were not optimally stained after GAF-fixation. RNAscope is an RNA *in situ* hybridization targeting single specific RNA molecules in individual cells in a variety of sample types including NBF-fixed paraffin-embedded tissues, representing a highly specific technique alternative to protein analysis in many human diseases.^8,9,70,71^ Our results on GAF-fixed samples were similar to what already found by RNAScope c-kit analysis on canine NBF-fixed mast cell tumors further indicating RNA preservation despite the mast cells loss on staining after GAF-fixation.^7^ On top of these results, GAF has the precious value of a very low toxicity.^22^ Glyoxal, despite being an irritating agent to skin and eyes, is not classified as a carcinogen, it has been found toxic only after long-term exposure by oral route and can be also used outside chemical hoods. ^5,17,26,31,36,48^

In conclusion, GAF is certainly deserving further attention from human and veterinary pathologists. Additional GAF applications should be conducted in teaching, research, and diagnostic for a wider recognition of GAF qualities and for reducing NBF drawbacks particularly concerning public health. In addition, scaling up GAF employment would conceivably reduce its costs which are not currently competitive with NBF. The present study validates GAF as a reliable encouraging fixative warranting special attention.

## Acknowledgements

We would like to acknowledge Dr. Cinzia Centelleghe for the technical support and Dr. Silvia Ferro for supporting in image editing.

## Authors’ contributions

FT, CG, LM, EM, EF, AR, FC, GDR, DD and LR designed and performed the experiments; VZ, PD, BB contributed to the experimental design; FT, VZ, AR, CG, MR, and SI performed histologic evaluations; VM performed statistical analysis; the manuscript was written by VZ and VM with contribution from the other authors.

